# Lack of functional caveolae in Cav3 mutated human dystrophic myotubes results in deficient mechanoprotection and IL6/STAT3 mechanosignaling

**DOI:** 10.1101/281113

**Authors:** Melissa Dewulf, Darius Köster, Bidisha Sinha, Christine Viaris de Lesegno, Valérie Chambon, Anne Bigot, Nicolas Tardif, Ludger Johannes, Pierre Nassoy, Gillian Butler-Browne, Christophe Lamaze, Cedric M. Blouin

**Affiliations:** Institut Curie – Centre de recherche, PSL Research University, Membrane dynamics and mechanics of intracellular signaling laboratory, CNRS UMR3666, INSERM U1143, Paris, France.; Centre for mechanochemical cell biology, University of Warwick, Coventry, United Kingdom.; Department of Biological Sciences, Indian Institute of Science Education and Research (IISER) Kolkata, Mohanpur, West Bengal, India.; Institut Curie – centre de recherche, PSL Research University, Endocytic trafficking and intracellular delivery laboratory, CNRS UMR3666, INSERM U1143, Paris, France.; Institut de Myologie, Sorbonne Universités, UPMC Université Paris 06, Thérapie des muscles striés laboratory, INSERM UMRS974, CNRS FRE3617, Paris, France.; LP2N, CNRS UMR 5298, IOA, Institut d’Optique Graduate School, Université de Bordeaux, Talence, France.

## Abstract

Caveolin-3 is the major structural protein of caveolae in muscle cells. Mutations in the *CAV3* gene cause different type of muscle disorders mostly characterized by defects in membrane integrity and repair, deregulation in the expression of various muscle proteins and deregulation of several muscle associated signaling pathways. We show here that myotubes derived from patients bearing the *CAV3* P28L and R26Q mutations present a lack of functional caveolae at the plasma membrane which results in an abnormal mechanoresponse. Mutant myotubes can no longer buffer the increase of membrane tension induced by mechanical stress and present an hyperactivation of the IL6/STAT3 signaling pathway at rest and under mechanical stress. The impaired mechanical regulation of the IL6/STAT3 signaling pathway by caveolae leads to chronic activation and a higher expression of muscle specific genes. These defects could be reversed by reassembling a pool of functional caveolae through expression of wild type Cav3. Our findings bring more mechanistic insight into human Cav3 associated muscle disorders and show a general defect in the mechanoresponse of *CAV3* P28L and R26Q myotubes.

## Introduction

Caveolae are cup-shaped plasma membrane invaginations that were first observed in the 50’s by Palade and Yamada on electron micrographs from vascular and gall bladder tissues (Palade, 1953, Yamada, 1955). Caveolae present a specific protein signature involving two main families of proteins, caveolins (caveolin-1, −2 and −3), and cavins (cavin-1, −2, −3 and −4) (Aboulaich et al., 2004; Hansen, Howard, and Nichols, 2011; Hill et al., 2008; Morén et al., 2012; Rothberg et al., 1992; Scherer et al., 1996; Way and Parton, 1995). Caveolins and cavins are expressed in almost every type of cells, except for caveolin-3 (Cav3) and cavin-4, which are expressed only in smooth and striated muscle cells (Way and Parton, 1995; Tagawa et al., 2008). Similarly to Cav1 in non muscle cells, Cav3 is necessary for the formation of caveolae at the plasma membrane of muscle cells (Minetti et al., 2002).

Caveolae have long been associated with several important cellular functions including endocytosis, lipid metabolism and cell signaling albeit with several unresolved controversies (Cheng and Nichols, 2016; Lamaze et al., 2017). More recently, a new function of caveolae was established as mechanosensors that play an essential role in cell mechanoprotection both *in vitro* and *in vivo* (Sinha et al., 2011; Lo et al., 2015; Garcia et al., 2017; Y.-W. Lim et al., 2017). The mechanical function of caveolae is likely to explain their particular abundance at the surface of many specialized cells that undergo chronic mechanical stress during their lifetime such as adipocytes, endothelial and muscle cells. Moreover, mutations or abnormal expression of caveolae components have been associated with lipodystrophy, vascular dysfunction, cancer and muscle disorders (reviewed in Lamaze et al., 2017; Le Lay and Kurzchalia, 2005). The molecular mechanisms underlying caveolin-associated diseases remain poorly understood.

In this study, we investigated the mechanical role of caveolae in caveolinopathies, a family of muscle diseases involving mutations in the *CAV3* gene. Human caveolinopathies can affect both cardiac and skeletal muscle tissues, and encompass five distinct genetic disorders: Rippling Muscle Disease (RMD), Distal Myopathies (DM), HyperCKemia (HCK), Limb-Girdle Muscular Dystrophy 1C (LGMD-1C), and Familial Hypertrophic Cardiomyopathy (HCM) (Gazzerro et al., 2010)., These disorders share nevertheless common characteristics including mild muscle weakness, high levels of serum creatine kinase, variations in muscle fiber size and an increased number of central nuclei as observed in muscle biopsies (Minetti et al., 1998; Carbone et al., 2000; Betz et al., 2001; Carbone et al., 2000; Tateyama et al., 2002).

We focused our investigations on two representative human caveolinopathies mutations namely Cav3 P28L and Cav3 R26Q, which are respectively responsible for HCK (Merlini et al., 2002), and RMD, HCK and LGMD-1C (Sotgia et al., 2003). These two heterozygous mutations both lead to the abnormal retention of Cav3 in the Golgi complex. The P28L and R26Q Cav3 mutations have so far been associated to deregulations of some signaling pathways (Sotgia et al., 2003; Brauers et al., 2010), defects in membrane repair (Hernandez-Deviez et al., 2008; Cai et al., 2009) and defect in the mechanoprotection of the muscle tissue (Lo et al., 2015). These studies were conducted either *in vivo* in transgenic mice or zebrafish or *in vitro* either in the mouse muscle cell line C2C12 or in fibroblasts overexpressing a mutant form of Cav3, and the mechanical function of caveolae was not extensively explored in this context.

In this study, we investigated whether the phenotype of human myotubes isolated from patients bearing the Cav3 P28L or Cav3 R26Q mutations could be associated with defects in the caveolae-mediated mechanoresponse operating at the plasma membrane. We also tested the new hypothesis that the mechanical flattening of caveolae could be involved in the selective regulation of some signaling pathways important for muscle physiology (Nassoy and Lamaze, 2012), by studying interleukin-6 (IL6) signaling, a pathway that plays an essential role in muscle tissue homeostasis (Muñoz-Cánoves et al., 2013).

We show here that the Cav3 P28L and Cav3 R26Q myotubes are unable to assemble *bona fide* caveolae at the plasma membrane, leading to a loss of membrane tension buffering and membrane integrity under mechanical stress. We also found that the absence of functional caveolae impairs the regulation of the IL6/STAT3 pathway in the mutant myotubes both at rest and under mechanical stress. As a result, the IL6 signaling pathway is chronically hyperactivated and the expression of several of the STAT3 target genes upregulated. Finally, the expression of WT Cav3 in mutant myotubes was sufficient to restore a functional pool of caveolae and to rescue mechanosensing and IL6/STAT3 signaling pathway regulation in these cells.

## Results

### Drastic decrease of caveolae number at the plasma membrane of Cav3 mutant myotubes

To address the impact of Cav3 mutations in human muscle disorders, we analyzed wild type (WT), Cav3 P28L and Cav3 R26Q myotubes obtained from immortalized myoblasts which were isolated from healthy or Cav3 mutant patients and differentiated for 4 days. The state of myotube differentiation was validated by the expression level of the differentiation marker MF-20 (myosin II heavy chain) in the three cell lines (Supplementary Fig. 1a). We first analyzed the presence and the ultrastructure of caveolae at the plasma membrane of myotubes by electron microscopy. In WT myotubes, we observed many structures corresponding to *bona fide* caveolae i.e. 60-80 nm cup-shaped invaginations that were directly connected to the plasma membrane or to larger vacuoles of variable size deeper inside the cell known as rosettes, and that could still be connected to the plasma membrane (Fig. 1a). In contrast, few caveolae if any could be detected at the plasma membrane of mutant myotubes, and no large vacuolar structures were observed.

**Figure 1.**
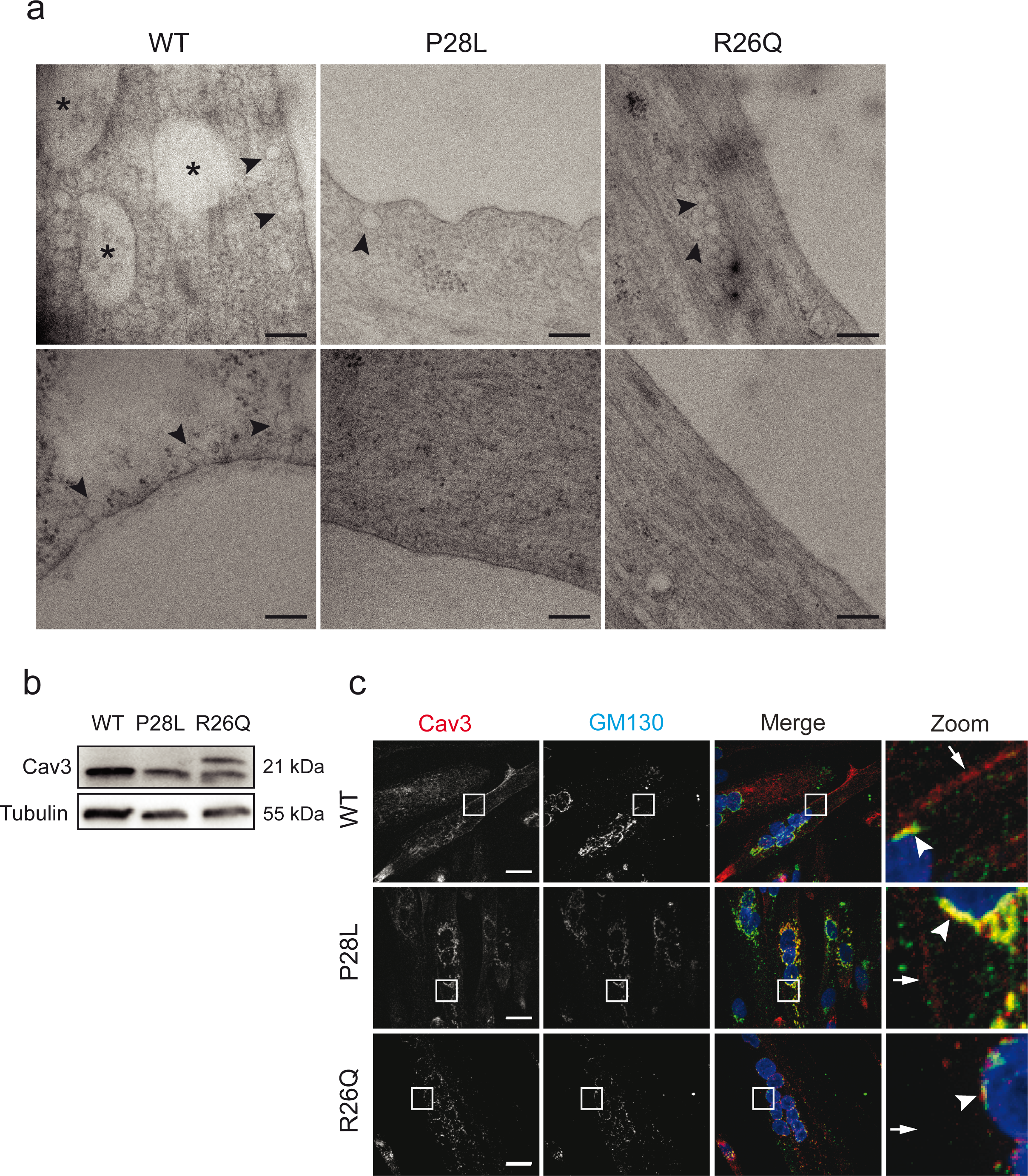
Characterization of caveolae and Cav3 expression in WT and Cav3 P28L and Cav3 R26Q myotubes. (**a**) Electron micrographs of WT, Cav3 P28L and Cav3 R26Q myotubes. Caveolae and interconnected caveolae are indicated with arrowheads and asterisks, respectively. (**b**) Immunoblot analysis of total levels of Cav3 in WT, Cav3 P28L and Cav3 R26Q differentiated myotubes. Tubulin serves as a loading control. (**c**) Immunofluorescent labeling of Cav3 and GM130 in WT, Cav3 P28L or Cav3 R26Q myotubes analyzed by confocal microscopy. Arrows in inset indicate the plasma membrane and arrowheads indicate the Golgi complex. (**a**) Scale bar = 200 nm. (**c**) Scale bar = 10 µm. Reproducibility of experiments: (**a**) Representative cells. (**b**) and (**c**) Representative data for 3 experiments.

This drastic decrease of caveolae number led us to investigate the localization of Cav3, since it is required for caveolae assembly at the plasma membrane (Minetti et al., 2002). Immunoblot analysis showed a reduced expression of mutant Cav3 as compared to WT (Fig. 1b), with a shifted band for the R26Q mutant corresponding to the Cav3 mutant form, as reported previously (Sotgia et al., 2003). Cav3 immunostaining revealed that WT Cav3 was mainly associated with the plasma membrane of myotubes and also partially localized in the Golgi complex, defined by GM130 staining (Fig. 1c). On the contrary, in the P28L and R26Q myotubes, Cav3 showed a drastic accumulation in the Golgi complex, as shown by the strong colocalization with GM130, in agreement with published data (Sotgia et al., 2003; Brauers et al., 2010). These data confirm that the Cav3 P28L and R26Q mutations retain Cav3 in the Golgi complex, which results in a drastic reduction of the number of caveolae present at the plasma membrane of the Cav3 mutant myotubes.

After 4 days of differentiation, Cav1 could still be expressed in myotubes and could potentially participate to the formation of caveolae independently from Cav3. We therefore analyzed Cav1 expression in myotubes after 4 days of differentiation and found that Cav1 was indeed expressed to the same level in the three cell lines (Supplementary Fig. 1b). We also found that Cav1 colocalized perfectly with Cav3 at the plasma membrane and to a lesser extent at the Golgi complex in WT myotubes whereas it was mainly retained in the Golgi complex in Cav3 P28L and R26Q myotubes (Supplementary Fig. 1c). This indicates that Cav1 can form hetero-oligomers with Cav3, and that the Cav3 P28L and R26Q mutants are dominant on Cav1 localization.

### Cav3 P28L and R26Q myotubes display major defects in membrane tension buffering and mechanoprotection under mechanical stress

To know whether the quasi absence of caveolae at the plasma membrane of mutated myotubes could induce defects in cell mechanoprotection, we first determined if the Cav3 P28L and R26Q myotubes could buffer the increase of membrane tension induced by mechanical stress. We thus applied a 45 mOsm hypo-osmotic shock to myotubes aligned by micropatterning and we measured the apparent membrane tension before and after 5 min of hypo-osmotic shock using membrane nanotube pulling with optical tweezers as described (Sinha et al., 2011). While the mutant myotubes showed no significant changes in membrane tension in resting condition (Fig. 2a), they showed a significant increase of membrane tension (P28L: 63.2% ± 7.47%; R26Q: 93.8% ± 11.2%) under 45 mOsm hypo-osmotic shock compared to WT myotubes (37.9% ± 8.93%) (Fig. 2b). These results clearly show that the Cav3 P28L and R26Q mutant myotubes have lost the ability to buffer membrane tension variations induced by intense mechanical stress.

**Figure 2.**
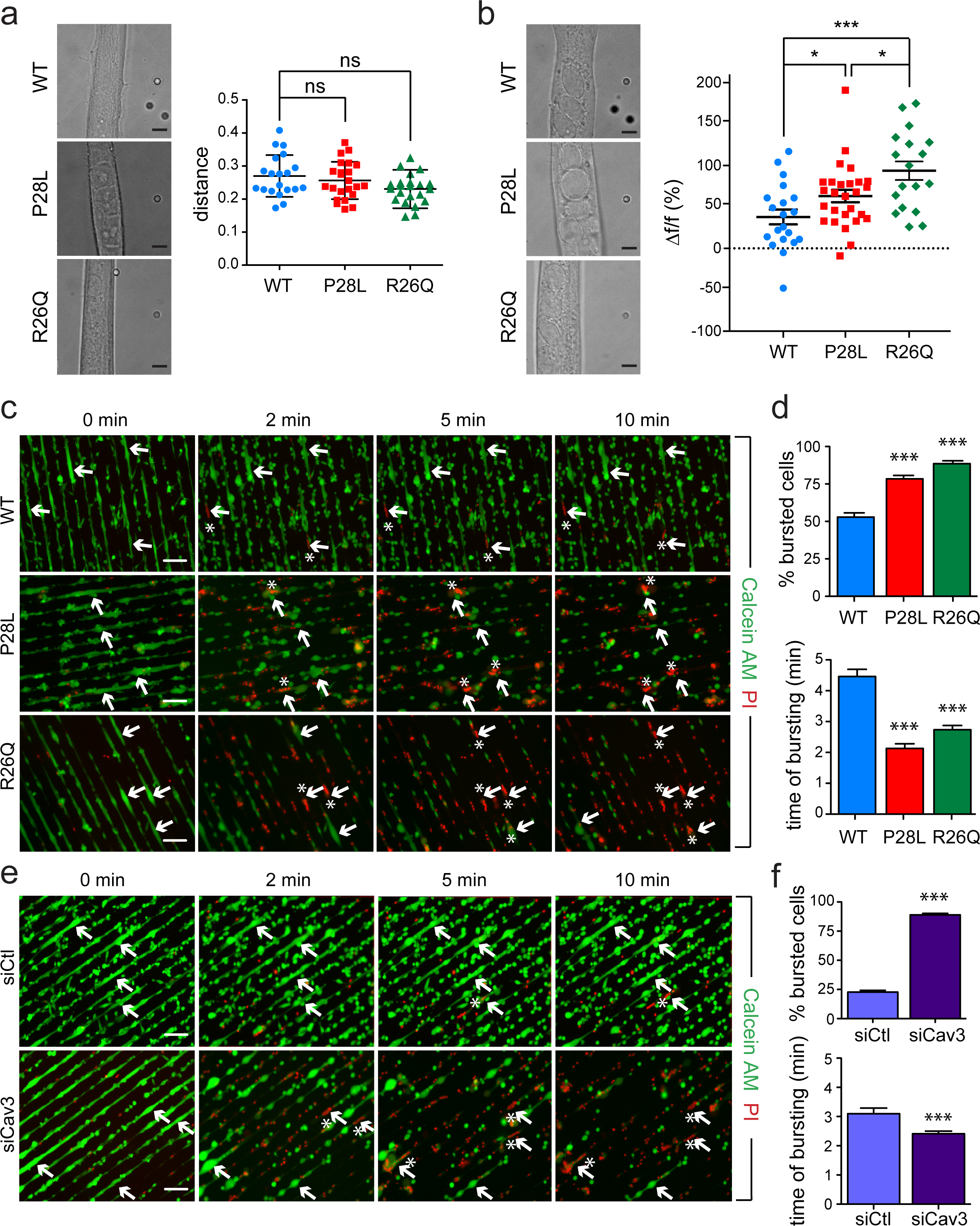
Cav3 P28L and Cav3 R26Q myotubes present defects in membrane tension buffering and membrane integrity. (**a, b**) Membrane tension measurements analysis using optical tweezers and nanotube pulling on micropatterned WT, Cav3 P28L or Cav3 R26Q myotubes. Membrane tethers were pulled in the perpendicular axis of aligned myotubes after micropatterning in resting conditions and 5 min after a 45 mOsm hypo-osmotic shock (**a, b**, left panels). Membrane tension was analyzed in resting condition (**a,** right panel) and the difference of membrane tension before and after hypo-osmotic shock was calculated, reflecting the percentage of increase of membrane tension upon mechanical stress (**b,** right panel). (**c, e**) Micropatterned WT, Cav3 P28L or Cav3 R26Q myotubes (**c**) WT ctl (siCtl) and Cav3-depleted (siCav3) myotubes (**e**) were loaded with calcein-AM (green). The medium was switched with a 30 mOsm medium supplemented with propidium iodide (PI, red). Representative pictures were taken at the indicated times during hypo-osmotic shock. Arrows correspond to myotubes and asterisks correspond to burst myotubes. (**d, f**) Quantification of the percentage of burst myotubes (upper panel) and mean time of bursting in minutes (lower panel) in (**c**) and (**e**), respectively. (**a, b**) Scale bar = 5 µm. (**c, e**) Scale bar = 120 µm. Reproducibility of experiments: (**a**) Representative pictures and quantifications from 7 independent experiments (WT n=20, P28L n=23 and R26Q n=22) (**b**) Representative pictures and quantifications from 7 independent experiments (WT n=20, P28L n=27 and R26Q n=18). (**c**) Representative data of 3 independent experiments quantified in (**d)** (% burst cells: WT n=310, P28L n=299 and R26Q n=271; mean time of bursting: WT n=165, P28L n=233 and R26Q n=240). (**e**) Representative data of 3 independent experiments quantified in (**f)** (% burst cells: siCtl n=749 and siCav3 n=569; mean time of bursting: siCtl n=171 and siCav3 n=506). Mean value ± SD. (**a**, **b**) Statistical analysis were done using Kurskal-Wallis test. (**d**, **f**) Statistical analysis with two-tailed unpaired t test; * P<0,05, *** P<0,001.

We next tested whether the lack of membrane tension buffering could result in insufficient mechanoprotection and increased membrane fragility in mechanically challenged mutant myotubes. We designed an assay to quantify the percent of cells that rupture their membrane under mechanical stress. This membrane bursting assay consists in incubating micropatterned myotubes with calcein-AM, a permeant green fluorescent dye that becomes fluorescent only inside the cell, and with the nucleus specific blue dye DAPI to specifically visualize myotubes by nuclei staining (Supplementary Fig. 2b). Live imaging was performed on myotubes subjected for 10 min to a 30 mOsm hypo-osmotic shock in the presence of propidium iodide (PI), a non-permeant red fluorescent dye that cannot enter cells if the plasma membrane is intact. The concomitant decrease of calcein-AM fluorescence and the appearance of PI fluorescence in the nucleus indicate a loss of membrane integrity (Fig. 2c). After 10 min of hypo-osmotic shock, in comparison to WT myotubes, mutant myotubes not only showed a higher percentage of cells that bursted out (WT: 52.9% ± 2.84%; P28L: 78.26% ± 2.38%; R26Q: 88.56% ± 1.93%) but also a shorter time of resistance to bursting (WT: 4.5 min ± 0.2, P28L: 2.1 min ±0.1, R26Q: 2.7 min ±0.2) (Fig. 2d). Importantly, when we apply a milder hypo-osmotic shock (150 mOsm), for which no increase in membrane tension could be measured, the plasma membrane of all three cell lines remained intact after 10 min of shock (Supplementary Fig. 2b and 2c).

In agreement with the absence of caveolae observed at the plasma membrane of the mutant myotubes (Fig. 1), when WT myotubes were depleted for Cav3, we measured a percentage of bursted out cells that was similar to mutant myotubes (siCtl: 22.7% ± 1.5%, siCav3: 88.93% ± 1.4%; Fig. 2e and 2f; Supplementary Fig. 2d). Likewise, Cav3 depleted myotubes significantly bursted out faster as compared to control myotubes (siCtl: 3.1 min ± 0.2, siCav3: 2.4 min ± 0.1) (Fig. 2e and 2f). Together, our results demonstrate that the Cav3 P28L and R26Q mutant myotubes are unable to provide the mechanoprotection that is required to maintain the integrity of the plasma membrane under mechanical stress and behave similarly to myotubes depleted for Cav3.

### Chronic hyperactivation of IL6/STAT3 signaling in Cav3 P28L and R26Q mutant myotubes

Considering the key role of caveolae and caveolin in intracellular signaling (Lamaze et al., 2017), we next investigated the possible impact of the loss of functional caveolae on some of the key signaling pathways in the muscle. We focused our analysis on the IL6/STAT3 signaling pathway that has been associated with satellite cells exhaustion and muscle wasting (Bonetto et al., 2011; Price et al., 2014; Tierney et al., 2014). Furthermore, the IL6 signal transducer glycoprotein gp130, which, together with the IL6 receptor subunit, assemble the IL6 receptor, has been localized in caveolae in a myeloma cell line (Podar et al., 2003), suggesting a potential regulation of the IL6 signaling pathway by caveolae. IL6 binding to the IL6 receptor is classically followed by the activation of receptor-bound JAK1 and JAK2 kinases, which in turn phosphorylate the signal transducer and activator of transcription 3 (STAT3) that is then translocated as a dimer to the nucleus where it activates the transcription of IL6 sensitive genes (Heinrich et al., 2003).

We therefore monitored the level of STAT3 activation i.e. tyrosine phosphorylation (pSTAT3) in myotubes stimulated for 5 and 15 min with physiological concentrations of IL6. At steady state, in the absence of IL6 stimulation, little if any tyrosine phosphorylation of STAT3 could be detected in WT myotubes. In contrast, we found that the level of pSTAT3 was already higher in Cav3 P28L and R26Q mutant myotubes even in the absence of IL6 stimulation. While IL6 stimulation for 5 and 15 min led to increased levels of pSTAT3 in WT myotubes, we observed a higher level of pSTAT3 in Cav3 P28L and R26Q mutant myotubes at similar times of stimulation (Fig. 3a, b). To rule out a possible contamination of cell lysates by undifferentiated myotubes, we also investigated the nuclear translocation of pSTAT3 by immunofluorescence directly in differentiated myotubes characterized by the presence of multiple nuclei (Fig. 3c, d). Again, we could detect a higher level of pSTAT3 in the nuclei of mutant myotubes as compared to WT at steady state (WT: 1.62 ± 0.1; P28L: 2.06 ± 0.2; R26Q: 2.02 ± 0.2), and after 15min of IL6 stimulation, although it was less pronounced in P28L mutants (WT: 3.03 ± 0.2; P28L: 3.66 ± 0.2; R26Q: 4.8 ± 0.4). These data confirm the immunoblot analysis, and show that the IL6/STAT3 signaling pathway is hyperactivated in the Cav3 P28L and R26Q mutant myotubes both at steady state and upon IL6 stimulation.

**Figure 3.**
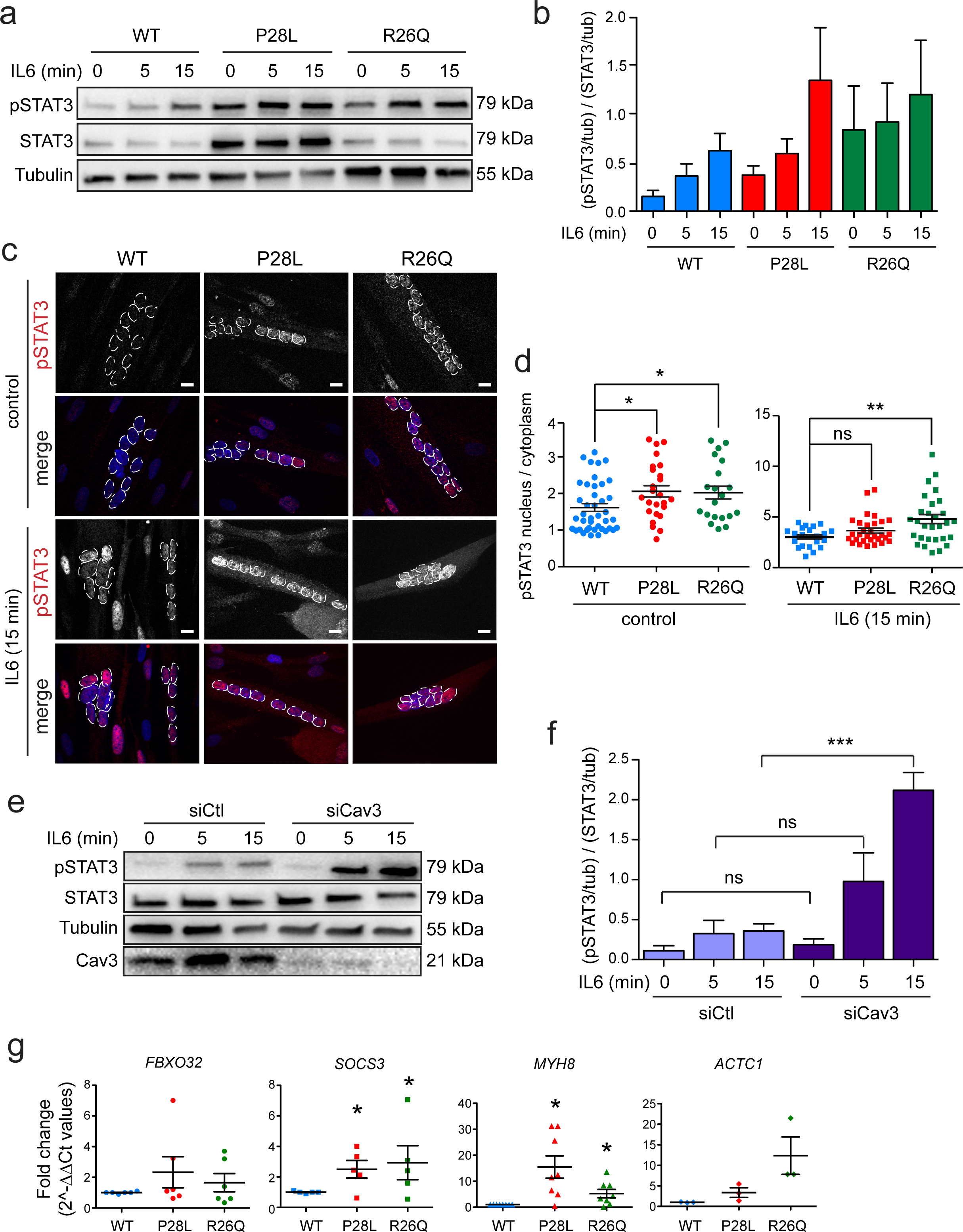
Hyperactivation of IL6/STAT3 signaling in Cav3 P28L and Cav3 R26Q myotubes. (**a**) Immunoblot analysis of pSTAT3 and STAT3 levels in WT, Cav3 P28L and Cav3 R26Q myotubes stimulated for the indicated times with 10 ng/mL IL6. Tubulin serves as a loading control. (**b**) Quantification of STAT3 activation of (**a**), corresponding to the ratio pSTAT3 on STAT3 total levels after normalization to tubulin levels. (**c**) Confocal microscopy of immunofluorescent pSTAT3 in WT, Cav3 P28L and Cav3 R26Q myotubes stimulated or not for 15 min with 10 ng/mL IL6. (**d**) Quantification of pSTAT3 nuclear translocation in (**c**) corresponding to nuclei/cytoplasm mean intensity ratio of pSTAT3. (**e**) Immunoblot analysis of pSTAT3 levels in WT ctl (siCtl) and Cav3-depleted (siCav3) myotubes stimulated for the indicated times with 10 ng/mL IL6. (**f**) Quantification of STAT3 activation in (**e**), corresponding to the ratio pSTAT3 on STAT3 total levels after normalization to tubulin levels. (**g**) Expression of STAT3 related genes: from left to right *FBXO32, SOCS3*, *MYH8* and *ACTC1* in WT, Cav3 P28L or Cav3 R26Q myotubes. (**c**) Scale bar = 10 µm. Reproducibility of experiments: (**a**, **c** and **e**) Representative data. (**b**) Quantification was done on 4 independent experiments. (**d**) Quantification was done on 3 independent experiments (0 min: WT n=41, P28L n=25, R26Q n=21; 15 min: WT n=22, P28L n=30, R26Q n=30). (**f**) Quantification was done on 4 experiments. (**g**) Quantification was done on 6 (*FBXO32*), 5 (*SOCS3*), 8 (*MYH8*) and 3 (*ACTC1*) independent experiments. Mean value ± SEM. (**b**, **f**) Statistical analysis with two-tailed paired t test. (**d, g**) Statistical analysis with two-tailed unpaired t test * P<0,05, ** P<0,01, *** P<0,001, ns non significant.

We next investigated if the regulation of the IL6/STAT3 pathway was dependent on the presence of functional caveolae at the plasma membrane and thus on the expression of Cav3. We therefore repeated the experiments on WT myotubes depleted for Cav3. By immunoblot analysis, we found that Cav3 depletion by siRNA led also to an hyperactivation of the IL6 pathway with an overall activation of STAT3 (Fig. 3e, f) (0 min: siCtl: 0.11 ± 0.1, siCav3: 0.2 ± 0.1; 5 min: siCtl: 0.33 ± 0.2, siCav3: 0.98 ± 0.4; 15 min: siCtl: 0.4 ± 0.1, siCav3: 2.12 ±0.2). These results indicate that Cav3 is a negative regulator the IL6/STAT3 pathway in myotubes and that depletion of Cav3 in WT myotubes reproduces a similar phenotype than the Cav3 mutants. It demonstrates that the absence of Cav3 and/or caveolae at the plasma membrane of mutant myotubes is responsible for the constitutive hyperactivation of the IL6/STAT3 signaling pathway.

STAT3 is a key transcription factor which controls the transcription of many downstream genes whose products mediate the pleiotropic effects of STAT3 in physiological and pathological contexts (Levy and Lee, 2002). We therefore examined the consequences of IL6/STAT3 chronic activation on gene expression. In the context of muscle diseases, we investigated the transcription of muscle-specific genes as STAT3 was suggested to be involved in their regulation (Bonetto et al., 2011). We focused our analysis on *FBXO32,* a gene known to be associated with muscle atrophy, and on the *ACTC1*, *MYH8* and *SOCS3* genes that are associated with muscle regeneration. *SOCS3* serves also as a positive control, as it is transcribed upon STAT3 activation and its gene product SOCS3 is a major actor in the negative regulation of the pathway (Heinrich et al., 2003). Using quantitative PCR, we found an increased transcription of *SOCS3*, *MYH8* and *ACTC1* genes, and little effect on the *FBXO32* gene (*SOCS3*: WT 1, P28L 2.5 ± 0.58, R26Q: 2.95 ± 1.1; *FBXO32*: WT 1, P28L 2.33 ± 1.01, R26Q: 1.65 ± 0.59; *MYH8*: WT 1, P28L 15,15 ± 4,3, R26Q 5,24 ± 1,6; *ACTC1*: WT 1, P28L 3,37 ± 1,18, R26Q 12,3 ± 4,56) (Fig. 3g). These data strongly suggest that the high level of pSTAT3 found in the Cav3 P28L and R26Q mutant myotubes is responsible of the deregulation of several genes that could be involved in the muscular pathological phenotype.

### Impaired IL6/STAT3 mechanosignaling in Cav3 P28L and R26Q mutant myotubes

Although caveolae and caveolins have long been associated with signaling (Cheng and Nichols, 2016; Lamaze et al., 2017), the integration of this function with their role in mechanical stress has not yet been reported. We have proposed the hypothesis that the mechano-dependent cycle of caveolae disassembly and reassembly could be tightly linked to the regulation of some signaling pathways by caveolae (Lamaze et al., 2017; Nassoy and Lamaze, 2012). We thus analyzed whether the regulation of the IL6/STAT3 pathway by caveolae could depend on mechanical stress. When myotubes were subjected to hypo-osmotic shock and stimulated with IL6, we observed a dramatic decrease of STAT3 activation (approx. 80%) in WT myotubes whereas no significant change was observed in Cav3 P28L and R26Q mutant myotubes (Fig. 4a, b). We also tested the effect of mechanical stretching on IL6/STAT3 signaling as this is more relevant to the nature of mechanical stress experienced by skeletal muscles during exercise. When we applied a 10% cyclic stretch at 0.5 Hz for 30 min to WT myotubes followed by IL6 stimulation, we also observed a drastic reduction of STAT3 activation, confirming that the IL6/STAT3 pathway is tightly regulated by mechanical cues (Supplementary Fig. 3). To know whether the mechanical regulation of IL6 signaling requires the presence of functional caveolae, we also applied an hypo-osmotic shock to WT myotubes depleted of Cav3. While no effect was observed at steady state, we found that STAT3 activation was slightly decreased by mechanical stress (approx. 20%) in WT myotubes when no changes were observed in Cav3 depleted myotubes (Fig. 4c-d. The poor adhesion of Cav3 P28L and R26Q mutant myotubes on the stretching membrane did not allow to perform cycling stretching. Nevertheless, these results confirm that the IL6 pathway is negatively regulated by mechanical stress in myotubes and that this regulation is lost in the absence of functional caveolae as shown in Cav3 P28L and R26Q mutant myotubes and in WT myotubes depleted for Cav3. It further confirms that the defects of mechanoprotection and IL6/STAT3 signaling regulation observed in the Cav3 P28L and R26Q mutant myotubes result from the lack of caveolae assembly at their plasma membrane.

**Figure 4.**
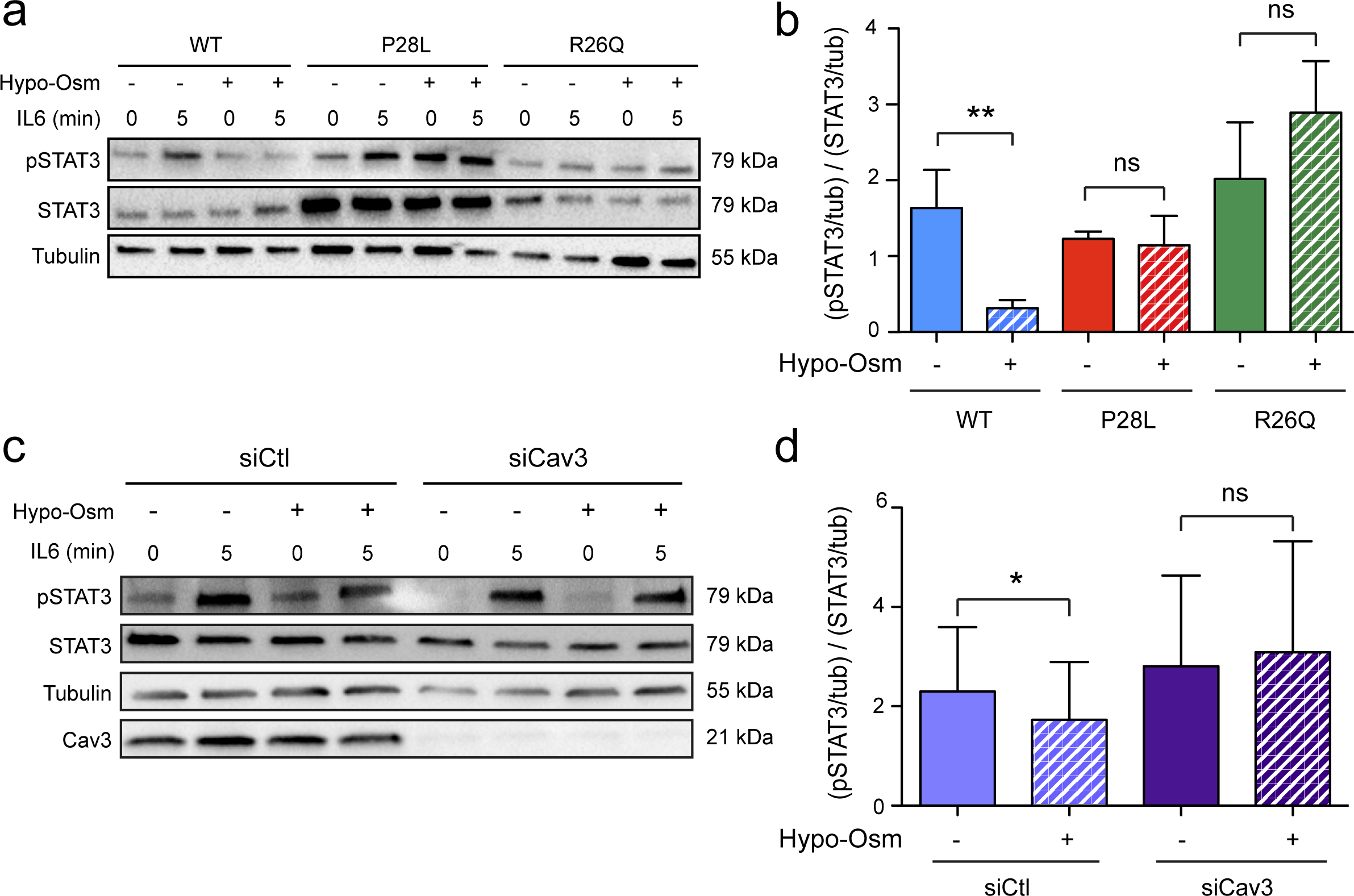
IL6/STAT3 mechanosignaling is impaired in Cav3 P28L and R26Q myotubes. (**a, c**) Immunoblot analysis of pSTAT3 and STAT3 levels in WT, Cav3 P28L and Cav3 R26Q myotubes. (**a**) WT ctl (siCtl) or Cav3-depleted (siCav3) myotubes (**c**) subjected or not to a 75 mOsm hypo-osmotic shock for 10 min, followed by stimulation or not with 10 ng/mL IL6 for 5 min. Tubulin serves as loading control. (**b, d**) Quantification of STAT3 activation in (**a**) and (**c**) respectively, corresponding to the ratio pSTAT3 on STAT3 total levels after normalization to tubulin levels. Reproducibility of experiments: (**a, c**) Representative data. (**b**) Quantification was done on 5 and 3 independent experiments for WT and mutants respectively. (**d**) Quantification was done on 4 independent experiments. Mean value ± SEM. (**b, d**) Statistical analysis with two-tailed paired t test, * P<0,05, ** P<0,01, ns = non significant.

### Expression of WT Cav3 is sufficient to rescue a normal phenotype in Cav3 P28L myotubes

Our experiments show that the depletion of Cav3 in WT myotubes reproduces the defects observed in P28L and R26Q myotubes, implying that the absence of Cav3 at the plasma membrane, as a result of its abnormal retention in the Golgi complex, is responsible for the observed phenotype. To validate this hypothesis, we generated stable WT and P28L myoblasts expressing either GFP or Cav3 WT tagged with GFP (Cav3-GFP). Immunofluorescent microscopy confirmed that re-expressed Cav3-GFP was mainly localized at the plasma membrane whereas endogenous Cav3 remained colocalized with the Golgi marker GM130 in Cav3-GFP P28L myotubes (Fig. 5a and supplementary Fig. 4a). We next performed electron microscopy to see if Cav3 WT expression would allow to reconstitute a pool of structurally defined caveolae at the plasma membrane of Cav3 P28L myotubes expressing GFP or Cav3 GFP. While the plasma membrane of control GFP myotubes presented very few, often isolated, caveolae structures, Cav3 rescued myotubes presented higher amount of *bona fide* caveolae, including larger vacuolar structures with connected caveolae i.e. rosettes, as observed above in WT myotubes (Fig. 5b and 1c). These observations confirm that the decrease of caveolae numbers in Cav3 P28L myotubes is a direct consequence of the retention of Cav3 P28L in the Golgi complex.

**Figure 5.**
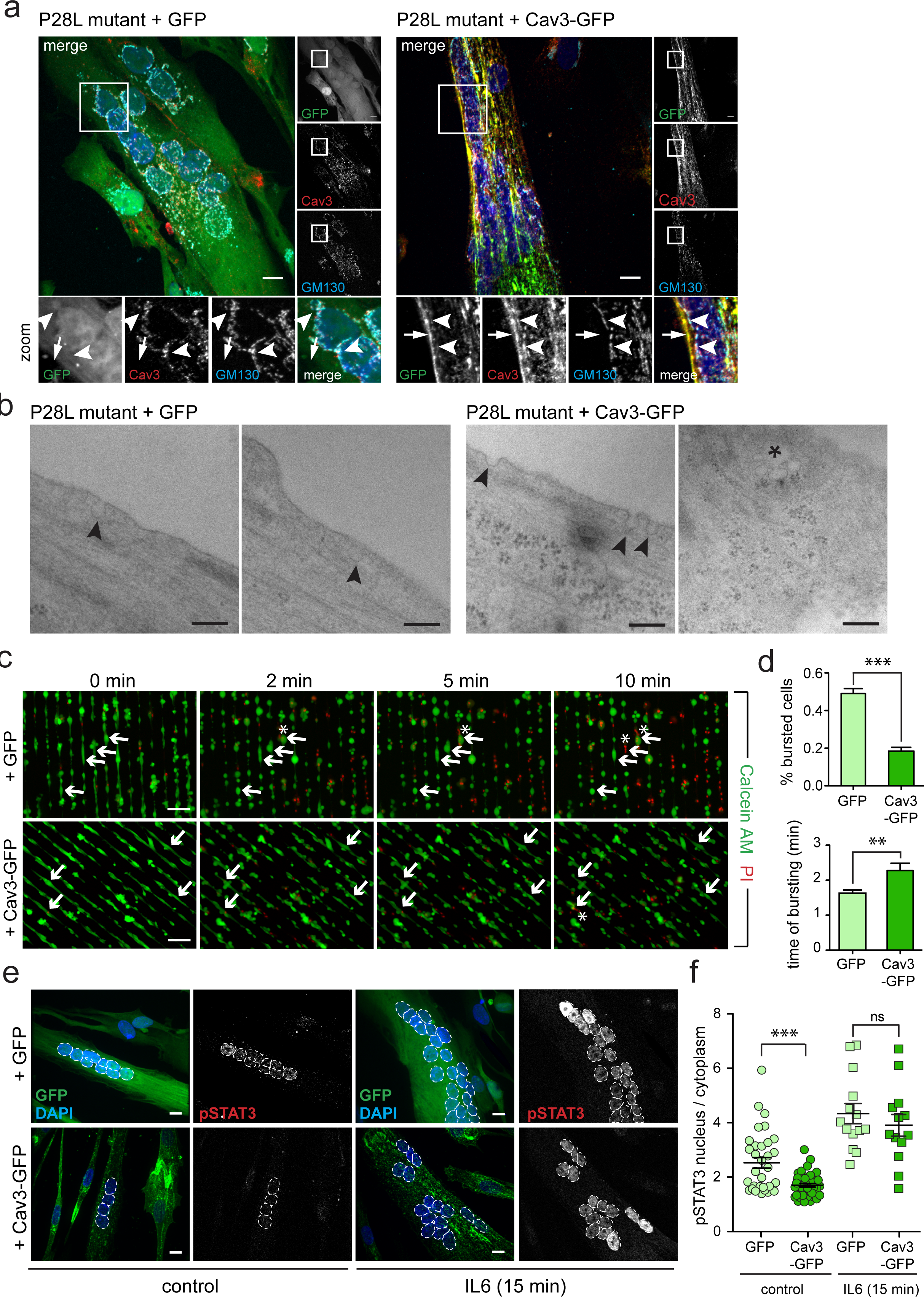
Expression of WT Cav3 rescues a normal phenotype in Cav3 P28L and R26Q myotubes. (**a**) Immunofluorescent labeling of Cav3 and Golgi marker GM130 in Cav3 P28L GFP and P28L Cav3-GFP transduced myotubes analyzed by confocal microscopy. Arrows in inset indicate the plasma membrane and arrowheads indicate the Golgi complex. (**b**) Electron micrographs of Cav3 P28L GFP and P28L Cav3-GFP transduced myotubes. Caveolae and interconnected caveolae are indicated with arrowheads and asterisks, respectively. (**c**) Micropatterned Cav3 P28L GFP and P28L Cav3-GFP transduced myotubes were loaded with calcein-AM (green). The medium was switched with a 30 mOsm medium supplemented with propidium iodide (PI, red). Representative pictures were taken at the indicated times during hypo-osmotic shock. Arrows correspond to myotubes and asterisks correspond to burst myotubes. (**d**) Quantification of the percentage of burst myotubes (upper panel) and mean time of bursting in minutes (lower panel) in (**c**). (**e**) Confocal microscopy of immunofluorescent pSTAT3 in Cav3 P28L GFP or P28L Cav3-GFP transduced myotubes stimulated or not for 15 min with 10 ng/mL IL6. (**f**) Quantification of pSTAT3 nuclear translocation in (**e**) corresponding to nuclei/cytoplasm mean intensity ratio of pSTAT3. (**a**) Scale bar = 10 µm. (**b**) Scale bar = 200 nm. (**c**) Scale bar = 120 µm. (**e**) Scale bar = 10 µm. Reproducibility of experiments: (**a**) Representative pictures of 3 experiments. (**b**) Representative pictures. (**c**) Show representative data of 3 experiments quantified in (**d**) (% burst cells: GFP n=353 and Cav3-GFP n=358; time of burst: GFP n=175 and Cav3-GFP n=65). (**e**) Show representative data of 3 experiments quantified in (**f**) (control: GFP n=33 and Cav3-GFP n=42; 15 min: GFP n=14 and Cav3-GFP n=13). Mean value ± SEM. (**d**) and (**f**) Statistical analysis with a two-tailed unpaired t test, ** P<0,01, *** P<0,0001, ns = non significant.

The next question was to know whether the reconstitution of the caveolae reservoir at the plasma membrane of Cav3 P28L myotubes was sufficient to rescue the mechanoprotection and mechanosignaling properties observed in WT myotubes. We therefore monitored the resistance to bursting of P28L GFP and P28L Cav3-GFP myotubes as described above. Notably, Cav3-GFP P28L myotubes showed a dramatic increase in the resistance to bursting under hypo-osmotic shock as compared to P28L GFP myotubes (P28L GFP: 49.01% ± 2.7%; P28L Cav3-GFP: 18.44% ± 2.1%). It also took a significantly longer time for P28L Cav3-GFP myotubes to burst out as compared to P28L GFP myotubes (GFP: 1.63 min ± 0.09; Cav3-GFP: 2.28 min ± 0.21) (Fig. 5c, d). Finally, we analyzed the regulation of the IL6/STAT3 pathway by monitoring STAT3 phosphorylation and nuclear translocation in P28L GFP and P28L Cav3-GFP myotubes. At steady state, we observed a significant decrease of pSTAT3 activation and nuclear translocation in Cav3-GFP myotubes as compared to GFP myotubes, indicating that the expression of Cav3 was sufficient to reduce the hyperactivation of STAT3 observed at steady state in Cav3 P28L myotubes (P28L GFP: 2.53 ± 0.2, Cav3-GFP: 1.7 ± 0.1) (Fig 5e, f).

Interestingly, we also found that the expression of Cav3-GFP in WT myotubes led to a decrease of pSTAT3 activation and nuclear translocation at steady state as compared to non transduced WT myotubes (Supplementary Fig. 4c). Altogether, these results show that Cav3 expression and localization at the plasma membrane of P28L myotubes was sufficient to rescue the lack of membrane integrity and the hyperactivation of the IL6 pathway described above. We can therefore conclude that the phenotype observed in mutant myotubes is due to a lack of Cav3 and functional caveolae at the plasma membrane.

## Discussion

The first *CAV3* mutations associated with muscle disorders were described 20 years ago (Minetti et al., 1998). Since then, a lot of effort has been put in trying to understand how these mutations could alter muscle tissue integrity and cause the observed syndromes. Due to its localization at the plasma membrane of myotubes and at the sarcolemma of muscle fibers, Cav3 has been shown to regulate myoblasts fusion (Bjerregard et al., 2014) and T-tubules organization (Ferruccio Galbiati et al., 2001). Also, Cav3 interacts with the dystrophin complex (Song et al., 1996) and regulates the trafficking of dysferlin (Hernández-Deviez et al., 2006.), two important muscle proteins causing severe myopathies when deregulated. Furthermore, Cav3 is involved in the regulation of different signaling pathways important for muscle function such as calcium homeostasis (Weiss et al., 2008), the insulin/GLUT4/Akt pathway (Fecchi et al., 2006) or TrkA and EGFR signaling (Brauers et al., 2010). Finally, we showed a role of Cav3 in membrane tension buffering and mechanoprotection in P28L FHCK human myotubes (Sinha et al., 2011). Furthermore, a recent *in vivo* study showed in a model of zebrafish expressing the R26Q mutation, that Cav3 was required for the membrane integrity of muscle tissue (Lo et al., 2015).

In the present work, we studied human myotubes after differentiation from myoblasts that were isolated and immortalized from healthy patients or patients bearing either the Cav3 P28L or Cav3 R26Q mutation. As Cav3 and caveolae are likely to regulate several functions in myotubes, we chose to focus on two major but poorly explored functions of caveolae that could have an important impact on the integrity of muscle cells and tissue. We first studied mechanoprotection as this function recently described has not been extensively investigated in muscle cells and never in human myotubes. We demonstrate in the present study that the Cav3 P28L and R26Q myotubes show a major defect in mechanoprotection, with less resistance of the plasma membrane to mechanical stress. We show that this defect results from the absence of a functional reservoir of caveolae at the plasma membrane that leads to a lack of membrane tension buffering under mechanical stress.

We then studied the regulation of the IL6/STAT3 signaling pathway in myotubes at rest and under mechanical stress. Indeed, we have proposed that the cycles of caveolae disassembly and reassembly induced by mechanical stress could have an impact on some signaling pathways (Nassoy and Lamaze, 2012). We tested our hypothesis by investigating the IL6 pathway as it is tightly associated to mechanical stress in muscle cells, with IL6 being secreted mostly during exercise (Ostrowski et al., 1998). We could show for the first time that the Cav3 mutant myotubes present a major deregulation of the IL6/STAT3 signaling pathway, as this pathway was hyperactivated under resting conditions. This defect translated into increased gene expression of *MYH8*, *SOCS3* and *ACTC1*, genes that have been associated with muscle regeneration, a process that may be important for the regulation of muscle tissue size. In addition, we show that the IL6 signaling pathway is negatively regulated by mechanical stress in a Cav3 dependent manner, and that this mechano-regulation is lost in mutated myotubes. It is likely that the caveolae-dependent mechano-regulation of the IL6/STAT3 pathway allows a negative feedback loop to avoid a chronic activation of the pathway and to adapt it to the mechanical stress that myotubes constantly experience during their lifetime.

Importantly, we also found that Cav3 depleted myotubes phenocopy Cav3 mutant myotubes. Furthermore, the defects in mechanoprotection and IL6 signaling regulation could be, at least partially, rescued in Cav3 P28L myotubes re-expressing the WT form of Cav3 and thereby reassembling a reservoir of functional caveolae at the plasma membrane. In conclusion, our data confirm that the retention of the Cav3 mutated forms in the Golgi complex is responsible for the absence of functional and morphologically defined caveolae at the plasma membrane of mutated myotubes. This absence leads both to deficient mechanoprotection and impaired mechano-regulation of the IL6/STAT3 signaling pathway, a pathway tightly linked to muscle mass and size. Mechanoprotection and proper regulation of muscle size are two features extremely important for general muscle integrity and it is likely that the alteration of these two processes can be deleterious for muscle tissue integrity, as it is often seen in muscular dystrophic patients.

## Methods

### Cell lines

P28L and R26Q human myoblasts were immortalized by the platform for immortalization of human cells of the Institute of Myology as described in (Mamchaoui et al., 2011). Briefly, myoblasts were transduced with lentiviral vectors encoding hTERT and cdk4 and containing puromycin (P28L) or puromycin and neomycin (R26Q) selection markers. Transduced cells were selected with puromycin (1 µg/ml) for 6 days (P28L) or with puromycin (1 µg/ml) for 6 days and neomycin (1 mg/ml) for 10 days (R26Q). Cells were seeded at clonal density, and individual myogenic clones were isolated.

For caveolin-3 expression, immortalized WT and P28L myoblasts were transduced with lentiviral vectors expressing WT caveolin-3 and a GFP reporter gene (MOI 5). A GFP lentiviral vector was used as control (MOI 5).

### Cell culture

All cells were grown at 37°C under 5% of CO_2_. All myoblasts cell lines were cultured in Skeletal Muscle Cell Growth Medium (Promocell) supplemented with 20% FCS (Gibco, Life technologies), 50 µg/mL of fetuine, 10 ng/mL of epidermal growth factor, 1 ng/mL basic fibroblast growth factor, 10 µg/mL of insulin and 0,4 µ/mL of dexamethasone (Promocell). Prior to any cell seeding, surfaces (well, coverslip, patterned coverslips) are coated with 0.01% of matrigel (v/v) (Sigma) for 15 min at 37°C. For myoblast differentiation, confluent cells (80-100% confluency) are put in DMEM high-glucose Glutamax (Gibco, Life Technologies), supplemented with 0.1% of insulin (v/v) (Sigma) for 4 days.

### Antibodies and reagents

Mouse anti-αTubulin (Sigma-Aldrich, clone B512, T5168, 1/1000 for WB); mouse anti-caveolin-3 (Santa Cruz, clone A3, sc-5310, 1/1000 for WB, 1/250 for IF); rabbit anti-caveolin-1 (Cell Signaling, 3238, 1/1000 for WB, 1/500 for IF); goat anti-GM130 (Santa Cruz, clone P-20, sc-16268, 1/50 for IF); mouse anti-MF20 (kind gift of Vincent Mouly, 1/100 for WB, 1/20 for IF); mouse anti-STAT3 (Cell Signaling, clone 124H6, 9139, 1/1000 for WB); rabbit anti-pSTAT3 (Cell Signaling, clone D3A7, 9145, 1/1000 for WB, 1/75 for IF); Secondary antibodies conjugated to Alexa FITC, Cy3, Cy5 or horseradish peroxidase (Beckman Coulter or Invitrogen). DAPI (Sigma-Aldrich).

### RNA interference-mediated silencing

Myoblasts were transfected with small interfering RNAs (siRNAs) using HiPerFect (Qiagen) according to the manufacturer’s instructions at days 0 and 2 of differentiation and were cultured in differentiation medium for a total f 4 days. Experiments were performed on validation of silencing efficiency by immunoblot analysis using specific antibodies and normalizing to the total level of tubulin used as loading controls. 20 nM of a pool of four siRNA targeting Cav3 were used (SI03068730, SI02625665, SI02625658 and SI00146188, QIAGEN), Control siRNA (1022076, QIAGEN) was used at the same concentration and served as reference point.

### Immunoblotting

Cells were lysed in sample buffer (62.5 mM Tris/HCl pH 6.0, 2% v/v SDS, 10% glycerol v/v, 40mM dithiothreitol and 0.03% w/v phenol red). Lysates were analyzed by SDS–PAGE and Western blot analysis and immunoblotted with the indicated primary antibodies and horseradish peroxidase-conjugated secondary antibodies. Chemiluminescence signal was revealed using Pierce™ ECL Western Blotting Substrate, SuperSignal West Dura Extended Duration Substrate or SuperSignal West Femto Substrate (Thermo Scientific Life Technologies). Acquisition and quantification were performed with the ChemiDoc MP Imaging System (Bio-rad).

### Immunofluorescence

Myoblasts were grown and differentiated on coverslips for 4 days. For Cav3, Cav1, MF-20, GM130 staining, cells are fixed with 4% PFA (v/v) (Sigma-Aldrich) for 10min at RT, quenched in 50 mM NH_4_Cl and then permeabilized with 0.2% BSA (v/v) and 0.05% saponin (v/v) (Sigma-Aldrich) in PBS for 20 min. Cells are incubated sequentially with indicated primary and fluorescence-conjugated secondary antibody in permeabilization buffer for 1h at RT. For pSTAT3 staining, cells are fixed and permeabilized with cold methanol for 15 min at −20°C. After washes with PBS 0.2% BSA (v/v), cells are incubated sequentially with indicated primary and fluorescence-conjugated secondary antibody in PBS 0.2% (v/v) for 1h at RT. In both protocols, coverslips are mounted in Fluoromount-G mounting medium (eBioscience) supplemented with 2 µg/mL of DAPI (Sigma-Aldrich). Acquisition of images are done using a spinning disk microscope (inverted Spinning Disk Confocal Roper/Nikon; Camera: CCD 1392×1040 CoolSnap HQ2; objective : 60x CFI Plan Apo VC).

### Electron microscopy

Epon embedding was used to preserve the integrity of cell structures. Myotubes were fixed sequentially for 1 hour at room temperature with 1.25% glutaraldehyde in 0.1 M Na-Cacodylate and then overnight at 4°C.

Cells were washed extensively with 0.1 M Na-Cacodylate pH 7.2. Membrane fixation was performed for 1 h at room temperature with 1% OsO4 in 0.1 M Na-Cacodylate pH 7.2. Cells were dehydrated by incubation with aqueous solutions of ethanol at increasing concentrations (50, 70, 90, then 100%, each for 10 min at room temperature). Embedding was finally performed in LX112 resin. Cells were infiltrated with a 1:1 LX112:ethanol solution, washed with LX112, and embedded overnight at 60°C in LX112 resin. Ultrathin 65 nm sections were sliced using a Leica UCT ultramicrotome and mounted on nickel formvar/ carbon-coated grids for observations. Contrast was obtained by incubation of the sections for 10 min in 4% uranyl acetate followed by 1 min in lead citrate.

Electron micrographs were acquired on a Tecnai Spirit electron microscope (FEI, Eindhoven, The Netherlands) equipped with a 4k CCD camera (EMSIS GmbH, Münster, Germany)

### Micropatterning

18 mm coverslips were micropatterned as described in (Carpi et al., 2011) using a photo-mask with lines of 10 µm of width, separated by 60 µm. In both force measurements and membrane bursting assay, myoblasts are plated at confluency on line micropatterns coverslips coated with 0.01% of matrigel (v/v) (Sigma) for 15 min at 37°C. Differentiation of myoblasts is achieved as described above in section *Cell culture*.

### Force Measurements

Plasma membrane tethers were extracted from cells by a concanavalin A (Sigma-Aldrich) coated bead (3 µm in diameter, Polysciences) trapped in optical tweezers. The optical tweezers are made of a 1064 nm laser beam (ytterbium fiber laser, λ = 1064 nm, TEM 00, 5 W, IPG Photonics, Oxford, MA) expanded and steered (optics by Elliot Scientific, Harpenden, UK) in the back focal plane of the microscope objective (Apo-TIRF 100× NA 1.45, Nikon). The whole setup was mounted on a Nikon Eclipse-Ti inverted microscope. The sample was illuminated by transmitted light, and movies were acquired at 10 Hz with an EM-charge-coupled device camera (Andor iXon 897) driven by Micro-Manager. The fine movements and particularly the translational movement necessary to pull the membrane tether were performed using a custom-made stage mounted on a piezoelectric element (P753, Physik Instrumente, Karlsruhe, Germany) driven by a servo controller (E665, Physik Instrumente) and a function generator (Sony Tektronix AFG320).

Calibration was performed using an oscillatory modulation driven by a function generator and measuring the response of the bead to an oscillatory motion of the stage. We measured k = 159 pN/µm. This relationship is linear in the laser power range used for the experiments (0.4–1.2 W).

The membrane tether was held at constant length to measure the static force. For measuring membrane tension changes due to hypo-osmotic shock, a first tether was first pulled at 300 mOsm (iso condition). A second tube was pulled on the same cell 5 minutes after diluting the medium with milliQ water to obtain 45 mOsm. The position of the bead used to compute tether forces was detected from the images using a custom ImageJ macro.

### Membrane bursting assay

Line micropatterned myotubes are incubated in 5 µg/mL of calcein-AM (Life techonologies) and 50 µg/mL of DAPI (Sigma-Aldrich) for 15 min at 37°C in the dark. Medium was then switched back with differentiation medium to wash out the excess of calcein-AM. The medium is then switched again with a 30 mOsm hypo-osmotic shock medium obtained after a dilution of 10% medium and 90% H_2_O, supplemented with 2 mg/mL of PI (Sigma). Immediately after medium switching, pictures are taken every minute for 10 min using a videomicroscope (Inverted microscope Nikon Ti-E, Camera: CCD 1392×1040 CoolSnap HQ2, objective: 10x CFI Fluor).

### IL6 stimulation

Myotubes were starved 4h by switching the differentiation medium to DMEM medium. In resting conditions, cells are then stimulated by switching the medium with DMEM with 0.2% BSA (w/v), supplemented with 10 ng/mL of human recombinant IL6 (R&D) for 0, 5 or 15min at 37°C. For hypo-osmotic conditions, medium was first switched to 75% hypo-osmotic shock (25% DMEM, 75% H_2_O) for 5 min and then switched to the same medium supplemented with 10 ng/mL of IL6 for 5 more minutes at 37°C. For stretching conditions, myoblasts were differentiated on fibronectin (Sigma-Aldrich) coated stretchable plates (Uniflex® culture plate, Flexcell International) and were then subjected or not to 30 min of cyclic stretch (10% elongation, 0.5Hz), using the FX-4000T TM Tension Plus device (Flexcell International), followed or not by 5 min of IL6 stimulation. Cells are lysed and samples are analyzed by immunoblotting. For the analysis, pSTAT3 levels were quantified by calculating the ratio between pSTAT3 and STAT3, both normalized to Tubulin signal. For the analysis of pSTAT3 nuclear translocation, myotubes were differentiated on coverslips, stimulated with IL6 as described above for 0min or 15min and were then fixed for further immunofluorescence analysis. Quantification corresponds to the ratio between the mean pSTAT3 intensity in the nuclei on the one in the cytoplasm.

### Quantitative PCR

Cells were lysed and RNA extraction was performed using an extraction kit (RNeasy Plus, Qiagen). Reverse-transcription reaction was performed with 1 µg of RNA per reaction, using high capacity cDNA reverse-transcritpion kit (Applied biosystem). qPCR was performed on 50 ng of cDNA for a reaction in a total volume of 20 µL, using Taqman Gene Expression Assays (GAPDH: Hs02786624_g1; FBXO32: Hs01041408_m1; ACTC1: Hs01109515_m1; MYH8: Hs00267293_m1; SOCS3: Hs02330328_s1, Applied biosystem) and a Lightcycler 480 Probes Master kit (Roche). Relative expression levels were calculated using ΔΔCT method with fold changes calculated as 2^−ΔΔCT^.

### Statistical analyses

All analyses were performed using GraphPad Prism version 6.0 and 7.0, GraphPad Software, La Jolla California USA, www.graphpad.com. Two-tailed (paired or unpaired) t-test was used if comparing only two conditions. For comparing more than two conditions, Kurskal-Wallis test was used with Dunn’s multiple comparison test (if comparing all conditions to the control condition). Significance of mean comparison is marked on the graphs by asterisks. Error bars denote s.e.m or SD.

## Acknowledgments

We would like to thank Catherine Coirault and Stéphane Vassilopoulos, the laboratory of Bruno Goud and Nicolas Carpi for providing materials or expertise. We are grateful to Paolo Pierobon for help on the analysis of membrane tension measurements. The facilities as well as scientific and technical assistance from staff in the PICT-IBiSA/Nikon Imaging Centre at Institut Curie-CNRS and the France-BioImaging infrastructure (N°ANR-10-INSB-04) are acknowledged. The electron microscope facility was supported by the French National Research Agency through the “Investments for the Future” program (France-BioImaging, ANR-10-INSB-04). This work was supported by institutional grants from the Curie Institute, INSERM and CNRS, and by specific grants from Association Française contre les Myopathies (AFM): CAV-MUT (17151) to M.D; CAV-STRESS-MUS (14266 to C.M.B and 14293 to C.L)

## Author Contributions

M.D designed and performed the experiments, analyzed results and wrote the manuscript. D.K, B.S, C.V.L, V.C performed experiments or analysis. A.B, N.T, L.J, P.N, G.B.B provided technical support and conceptual advice. C.L supervised the project and designed experiments. C.M.B supervised the project, designed experiments and wrote the manuscript.

## Declaration

The authors declare no competing financial interest.

