## Supplementary Materials for "Lack of functional caveolae in Cav3 mutated human dystrophic myotubes results in deficient mechanoprotection and IL6/STAT3 mechanosignaling"

#### Supplementary figure legends

**Supplementary figure 1 | MF20 and Cav1 expression in WT, Cav3 P28L and R26Q myotubes.** (**a, b**) Immunoblot analysis of total levels of MF20 (**a**) and Cav1 (**b**) in WT, Cav3 P28L and Cav3 R26Q myotubes. Tubulin serves as a loading control. (**c**) Cav3, Cav1 and Golgi marker GM130 immunofluorescence were analyzed by confocal microscopy in WT, Cav3 P28L and Cav3 R26Q myotubes. (**a**), (**b**) and (**c**) Representative data for 3 experiments. (**c**) Scale bar = 10 µm.

**Supplementary figure 2 | Efficient membrane tension buffering and mechanoprotection in Cav3 P28L and Cav3 R26Q myotubes under mild hypo-osmotic shock.** (**a**) Calcein-AM and DAPI fluorescence of WT, Cav3 P28L and Cav3 R26Q myotubes prior to hypo-osmotic shock in the membrane bursting assay described in **Figure 2c**. Insets show DAPI in myotubes indicated with arrows in **Figure 2c**. (**b**) Micropatterned WT, Cav3 P28L and Cav3 R26Q myotubes were loaded with calcein-AM (green). The medium was switched with a 150 mOsm medium supplemented with propidium iodide (PI, red). Representative pictures were taken at the indicated times during hypo-osmotic shock. Arrows correspond to myotubes and asterisks correspond to burst myotubes. (**c**) Membrane tension measurements analysis using optical tweezers and nanotube pulling on micropatterned WT, Cav3 P28L and Cav3 R26Q myotubes. Membrane tethers were pulled in the perpendicular axis of myotubes after micropatterning in resting conditions and 5 min after a 150 mOsm hypo-osmotic shock (upper panel). Membrane tension was analyzed in resting condition (lower panel, left) and the difference of membrane tension before and after hypo-osmotic shock was calculated, reflecting the percentage of increase of membrane tension upon mechanical stress (lower panel, right) (**d**) Immunoblot analysis of Cav3 depletion in **Figure 2e**. (**a,b**) Scale bar = 120 µm. (**c**) Scale bar = 5µm Reproducibility of experiments: (**c**) Quantifications were done on 5 independents experiments (WT n=17, P28L n=16, R26Q n=14). Mean value ± SD. Statistical analysis were done using Kruskal-Wallis test, ns = non significant.

**Supplementary figure 3 | IL6/STAT3 signaling in WT myotubes under cyclic stretch.** (**a**) Immunoblot analysis of pSTAT3 and STAT3 levels in WT myotubes subjected or not to 30 min cyclic stretch. Myotubes were then stimulated or not with 10 ng/mL IL6 for 5 min. Tubulin serves as a loading control. (**b**) Quantification of STAT3 activation in (**a**) corresponding to the ratio pSTAT3 on STAT3 total levels after normalization to tubulin levels. Reproducibility of experiments: (**b**) Quantification was done on 3 experiments. Mean value ± SEM. Statistical analysis were done using two-tailed paired t test, * P<0,05.

**Supplementary figure 4 | Effect of Cav3 expression in mechanoprotection and IL6 signaling in WT myotubes.** (**a**) Immunofluorescent labeling of Cav3 and Golgi marker GM130 in WT GFP and WT Cav3-GFP transduced myotubes analyzed by confocal microscopy. Arrows and arrowheads in inset indicate the plasma membrane and the Golgi complex respectively. (**b**) Quantification of the percentage of burst myotubes after a 30 mOsm hypo-osmotic shock (left panel) and mean time of bursting in minutes (right panel) in (**a**). (**c**) Quantification of pSTAT3 nuclear translocation in WT GFP or WT Cav3-GFP transduced myotubes stimulated or not for 15 min with 10 ng/mL IL6, corresponding to nuclei/cytoplasm mean intensity ratio of pSTAT3. (**a**) Scale bar = 10 µm. Reproducibility of experiments: (**a**) Representative data from 3 independent experiments. (**b**) Quantification was done on 3 independent experiments (% burst: GFP n=714, Cav3-GFP n=610; time of burst: GFP n=80, Cav3-GFP n=171). (**c**) Quantification was done on 3 independent experiments (control: GFP n=16, Cav3-GFP n=21; 15 min: GFP n=16, Cav3-GFP n=21). (**b, c**) Statistical analysis were done using two-tailed unpaired t test, ***P<0,0001 *P<0,05, ns = non significant.

**Supplementary figure 1.** ***Dewulf et al.***

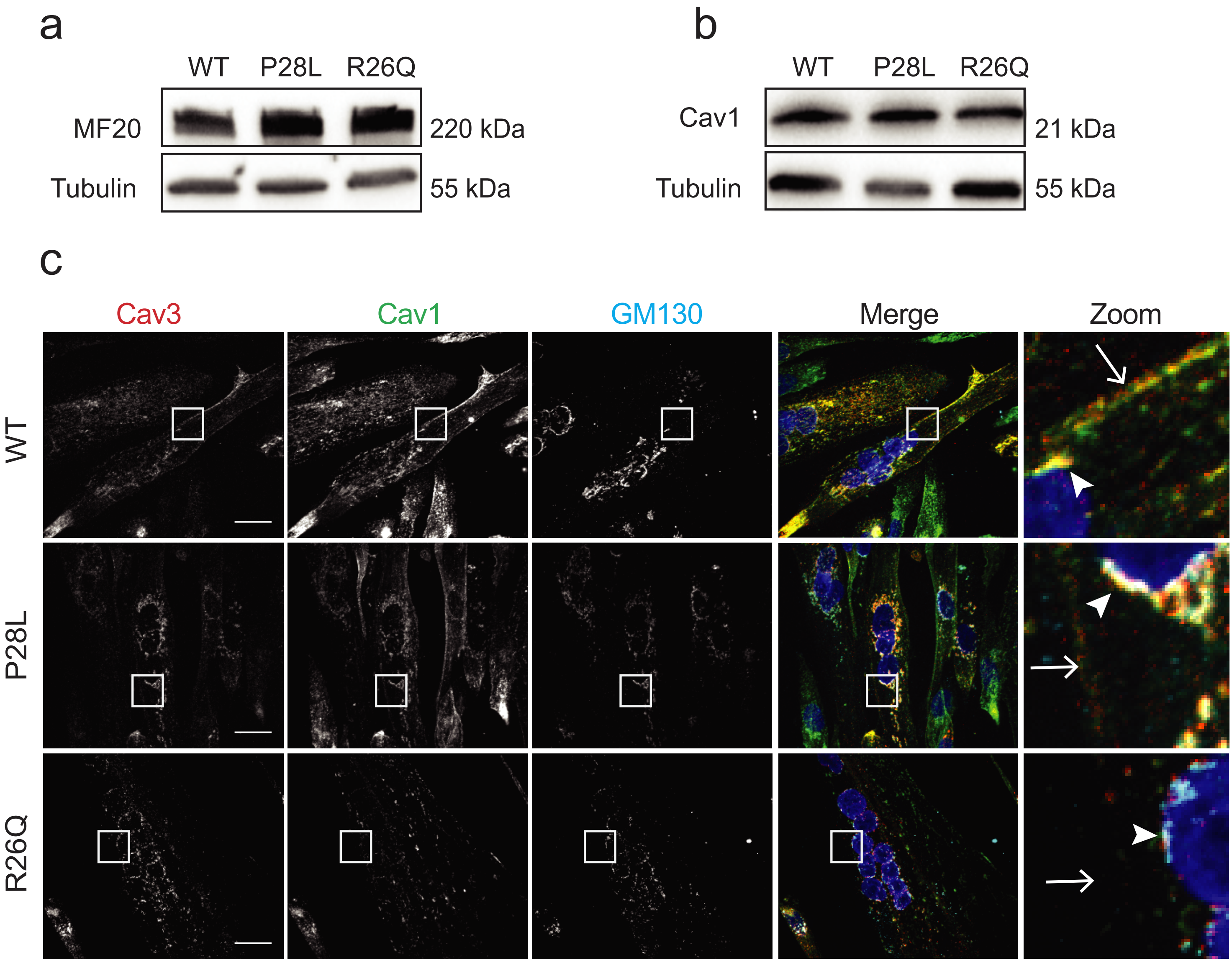

### Supplementary figure 2. *Dewulf et al.*

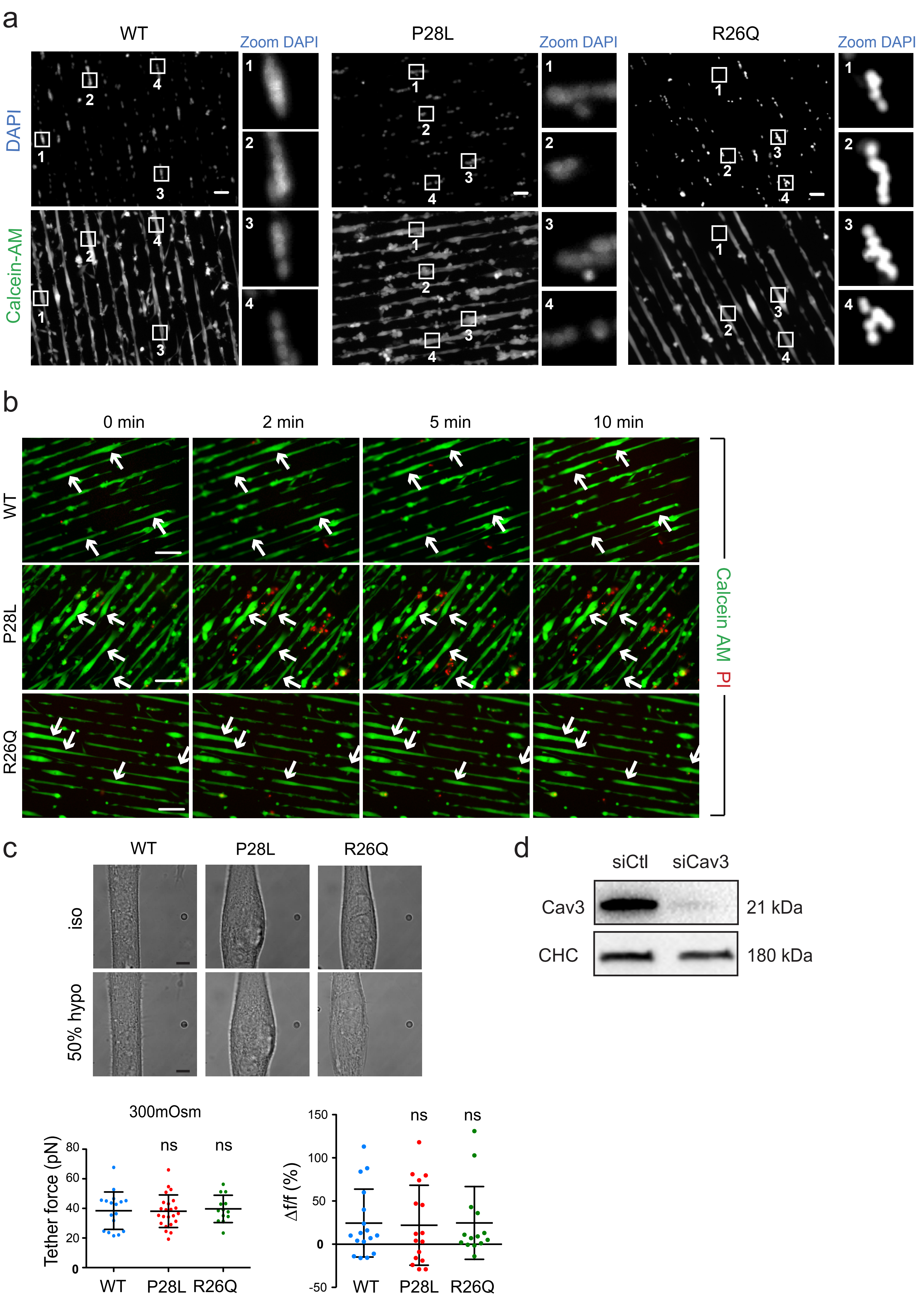

**Supplementary figure 3.** ***Dewulf et al.***

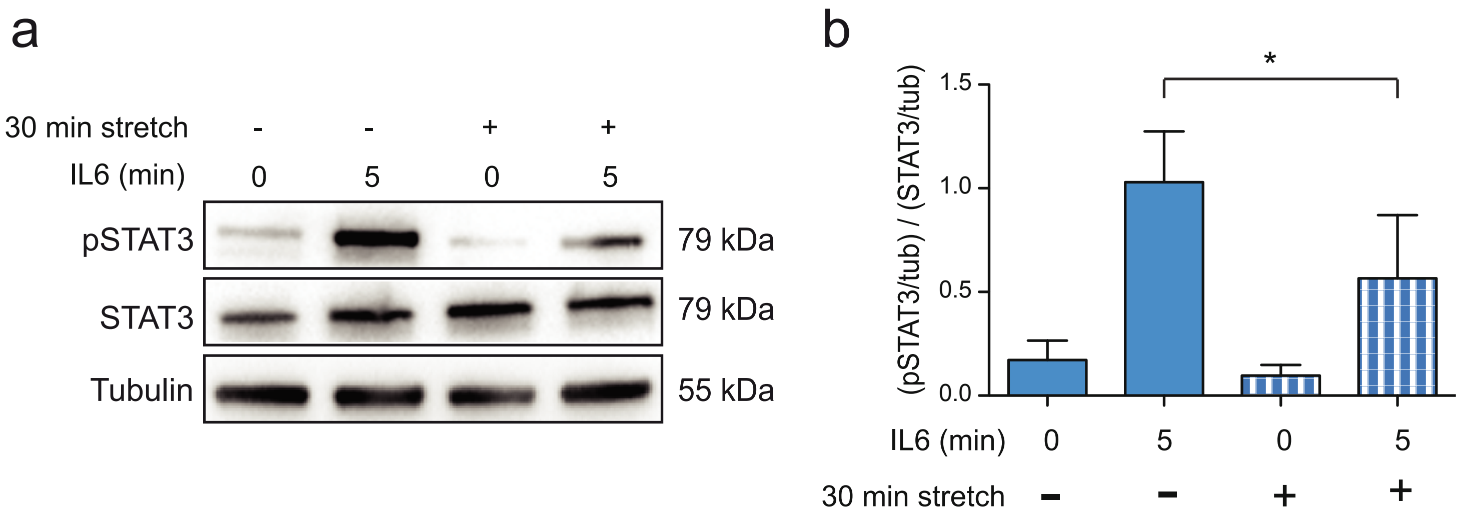

**Supplementary figure 4.** ***Dewulf et al.***

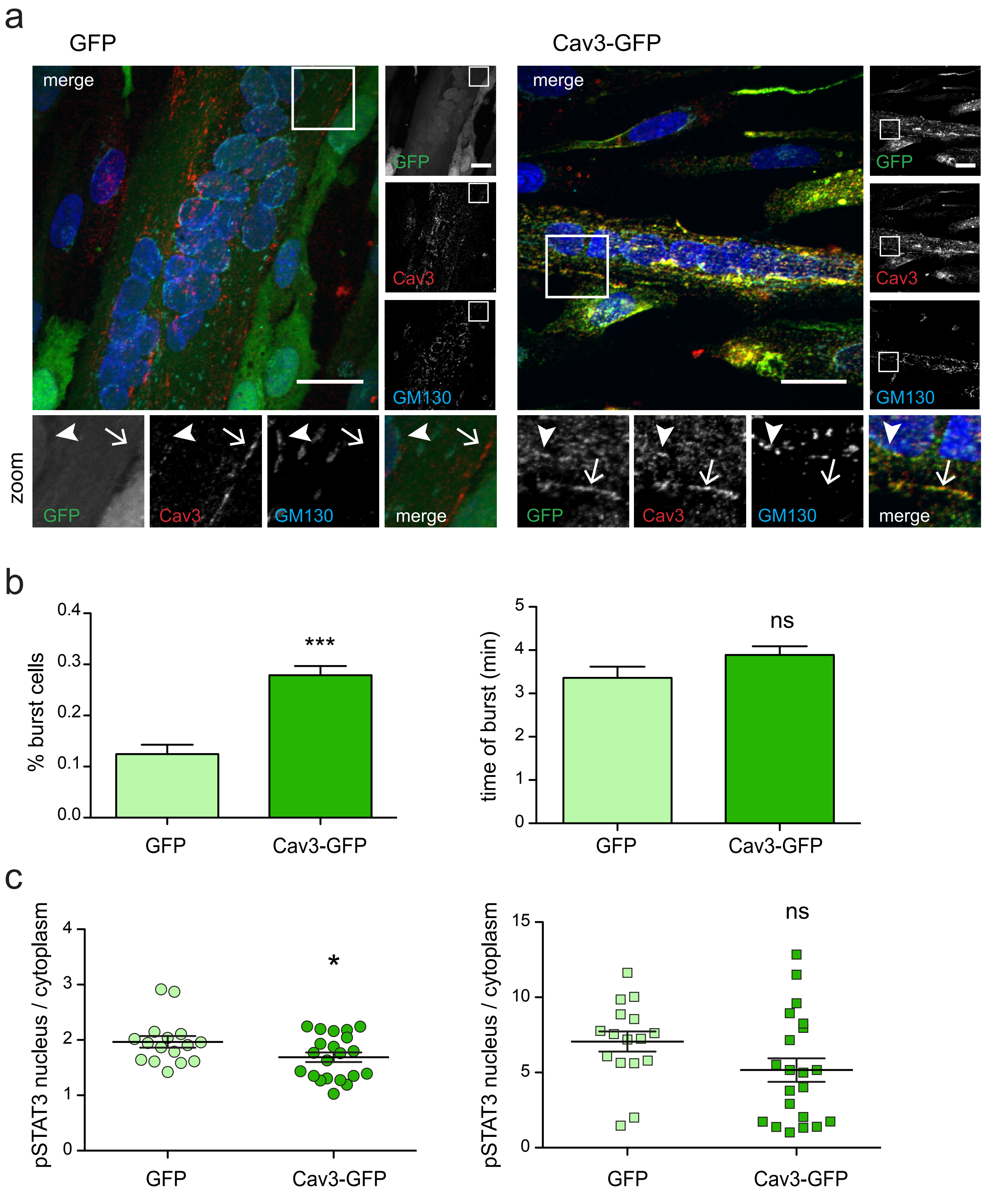
